# Policy complexity suppresses dopamine responses

**DOI:** 10.1101/2024.09.15.613150

**Authors:** Samuel J. Gershman, Armin Lak

## Abstract

Limits on information processing capacity impose limits on task performance. We show that animals achieve performance on a perceptual decision task that is near-optimal given their capacity limits, as measured by policy complexity (the mutual information between states and actions). This behavioral profile could be achieved by reinforcement learning with a penalty on high complexity policies, realized through modulation of dopaminergic learning signals. In support of this hypothesis, we find that policy complexity suppresses midbrain dopamine responses to reward outcomes, thereby reducing behavioral sensitivity to these outcomes. Our results suggest that policy compression shapes basic mechanisms of reinforcement learning in the brain.

## Introduction

Task performance is bounded by sensory and memory bottlenecks that limit the flow of information from task states to actions (Tishby and Polani, 2010; Gershman, 2020; Lai and Gershman, 2021). This implies that there is not a single performance optimum, but rather a spectrum of optima indexed by information capacity. This idea can be formalized by viewing an agent’s policy (the probabilistic mapping from states to actions) as a communication channel characterized by the mutual information between states and actions, also known as *policy complexity* (Figure 1). The channel’s capacity is an upper bound on policy complexity, which in turn determines an upper bound on achievable task performance.

**Figure 1:**
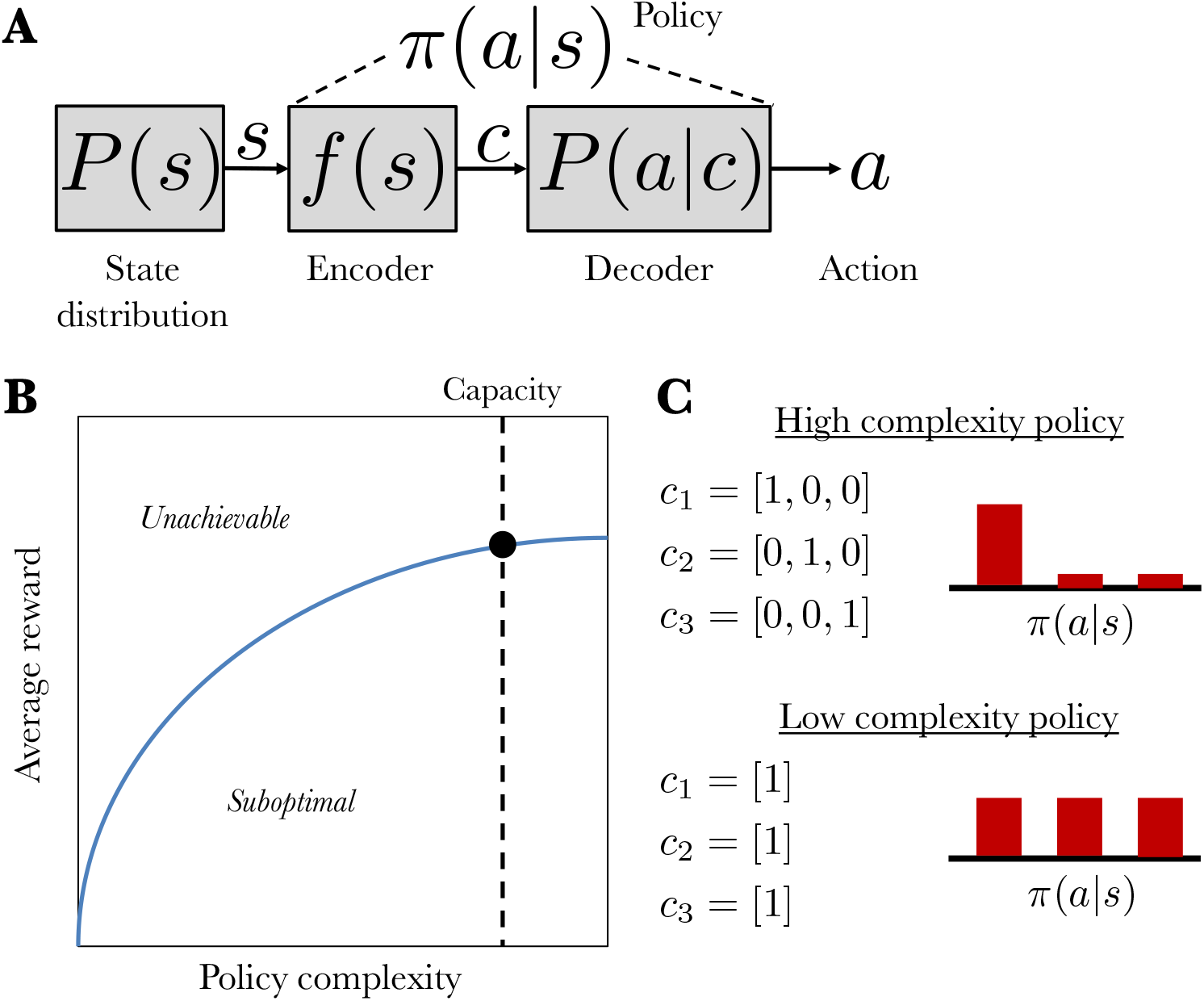
Policy compression framework. (A) State *s* is sampled from the state distribution and then compressed by an encoder compresses into codeword *c*. At the time of action selection, the codeword is probabilistically decoded into an action *a*. The complete mapping from states to actions is the policy, *π*(*a*|*s*). (B) The blue curve shows the optimal average reward achievable for each level of policy complexity (the mutual information between states and actions). A hypothetical capacity limit for an agent is shown as the dashed line; its intersection with the blue curve represents that agent’s maximum achievable average reward. All points above the blue line are unachievable, and all points below it are suboptimal. (C) Two example policies, distinguished by their complexity. The high complexity policy has a capacity of 3 bits and yields a low entropy distribution over actions. In contrast, the low complexity policy has a capacity of 1 bit and yields a high entropy distribution over actions.

Agents should *compress* their policies to stay within the capacity limit. Optimally compressed policies have several signatures: they are stochastic, biased towards high frequency actions, and sensitive to the distribution/number of states. These signatures have been documented experimentally in humans (Lai and Gershman, 2024), and have been argued to account for a wide range of well-established behavioral phenomena, such as perseveration (Gershman, 2020) and undermatching (Bari and Gershman, 2023).

Despite the abundance of behavioral evidence for policy compression, we still do not understand how it is implemented neurally. One possibility, motivated by reinforcement learning models of policy compression (Lai and Gershman, 2021, 2024), is that the reward prediction error signals used for policy updating register a policy complexity penalty, thereby driving policies towards a balance between reward maximization and policy compression. Specifically, the error δ should take the following form:

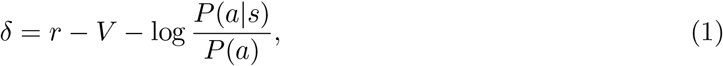

where *r* is the reward outcome, *V* is expected reward, and the last term is the policy cost (the probability of choosing action *a* in state *s* relative to the marginal probability of choosing *a* across all states), whose expectation is equal to policy complexity.

Since phasic dopamine signals classically conform to a reward prediction error signal (Schultz et al., 1997; Eshel et al., 2015; Gershman et al., 2024), we hypothesize that they will be suppressed by policy complexity. We tested this hypothesis using dopamine neuron recordings from mice during a perceptual decision task (Lak et al., 2020). The different stimuli in the task can be treated as distinct states (9 in total), allowing us to examine whether mice exhibit behavioral and neural signatures of policy compression.

## Materials and Methods

### Experimental procedure

The data analyzed in this paper were originally reported in Lak et al. (2020). We briefly summarize the data collection and analysis methods, referring readers to that paper for further details.

The data came from 5 mice (55 sessions total) performing a perceptual decision task (Fig. 2) while the activity of dopamine neurons in the ventral tegmental area were monitored using fiber photometry of GCaMP signals. Our analysis focused on the dopamine response at the time of reward feedback. Due to the relatively slow dynamics of the calcium signal, we averaged the signal between 300 and 800 ms following the outcome delivery, which encompasses the peak response. We then normalized the response by subtracting the calcium signal averaged over a 200 ms window centered on the outcome delivery. To aggregate across animals and sessions, we z-scored the responses within each session. We then fit a linear regression model to the responses with 4 regressors: an intercept, policy cost, action value (the average reward for the chosen action conditional on the current stimulus), and the outcome (water amount). For visualization of the policy cost effect, we calculated partial residuals (Larsen and McCleary, 1972), the differences between observed and predicted responses with the policy cost term removed; plotting this against policy cost isolates the cost regressor’s contribution after adjusting for the other regressors.

**Figure 2:**
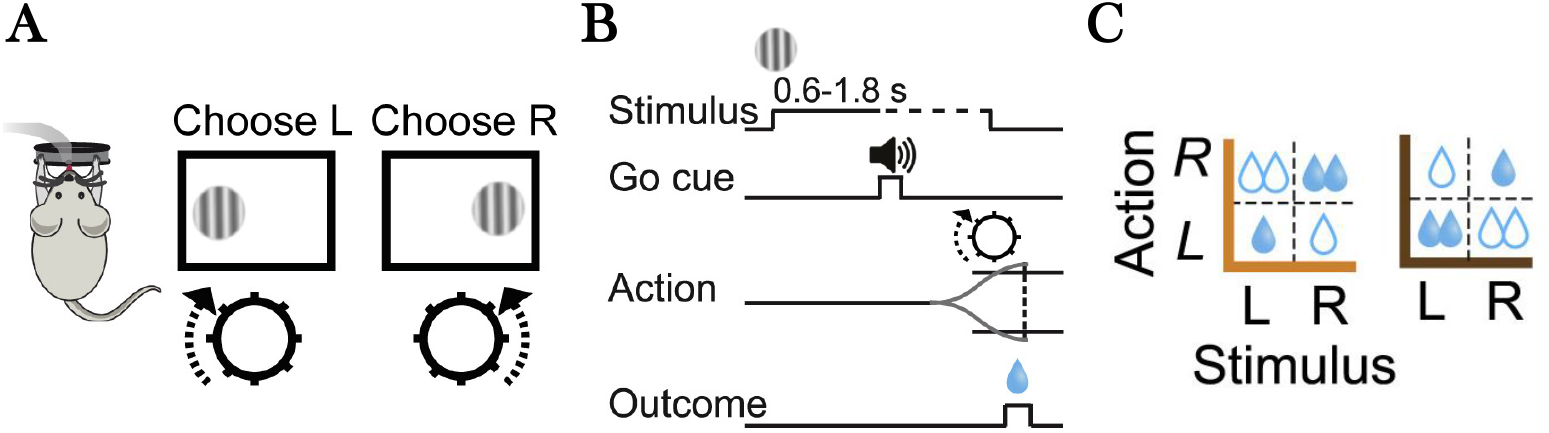
Task schematic. (A) Experimental interface. Mice reported the location (left or right) of a variable-contrast sinusoidal grating by turning a wheel. (B) Sequence of events on a single trial. Mice were required to await an auditory Go cue before responding, after which they received water reward for a correct action. (C) Reward structure. On different blocks, the correct action for one stimulus delivered twice as much reward as the correct action for the other stimulus.

### Policy compression model

Policy cost is defined as log *P* (*a*|*s*)*/P* (*a*), where *P* (*a*|*s*) is the conditional probability of action *a* given state *s* (here taken to be the stimulus), and *P* (*a*) is the marginal probability of action *a*. The probability distributions were estimated separately for each session. Policy complexity is defined as the average policy cost. An animal’s capacity is an upper bound on policy complexity. The optimal capacity-limited policy is given by *P* (*a*|*s*) ∝ exp[*βQ*(*s, a*) + log *P* (*a*)], where *Q*(*s, a*) is the average reward for taking action *a* in state *s*. The inverse temperature *β* is implicitly set based on the capacity limit and the task structure (see Lai and Gershman, 2021, for more details). We treated *β* as a free parameter, which we fit to the behavioral data using maximum likelihood estimation. To capture a small amount of state uncertainty, we smoothed the values across neighboring contrast levels.

## Results

On each trial, mice were presented with a sinusoidal grating on either the left or right side of the monitor, and had to report the side using a wheel following an auditory Go cue (Fig. 2A,B). Task difficulty was controlled by the contrast of the grating. In addition, the reward magnitude for correct actions was asymmetric across blocks of trials (Fig 2C).

We first checked for behavioral signatures of policy compression. We used the Blahut-Arimoto algorithm to calculate the optimal reward-complexity frontier (Fig. 3A). Each point on this frontier represents the maximum achievable average reward for a particular capacity limit. Points above the curve are unachievable, and points below the curve are suboptimal. We found that mouse behavior on this task was close to the optimal frontier, with a median deviation of 3.1%.

**Figure 3:**
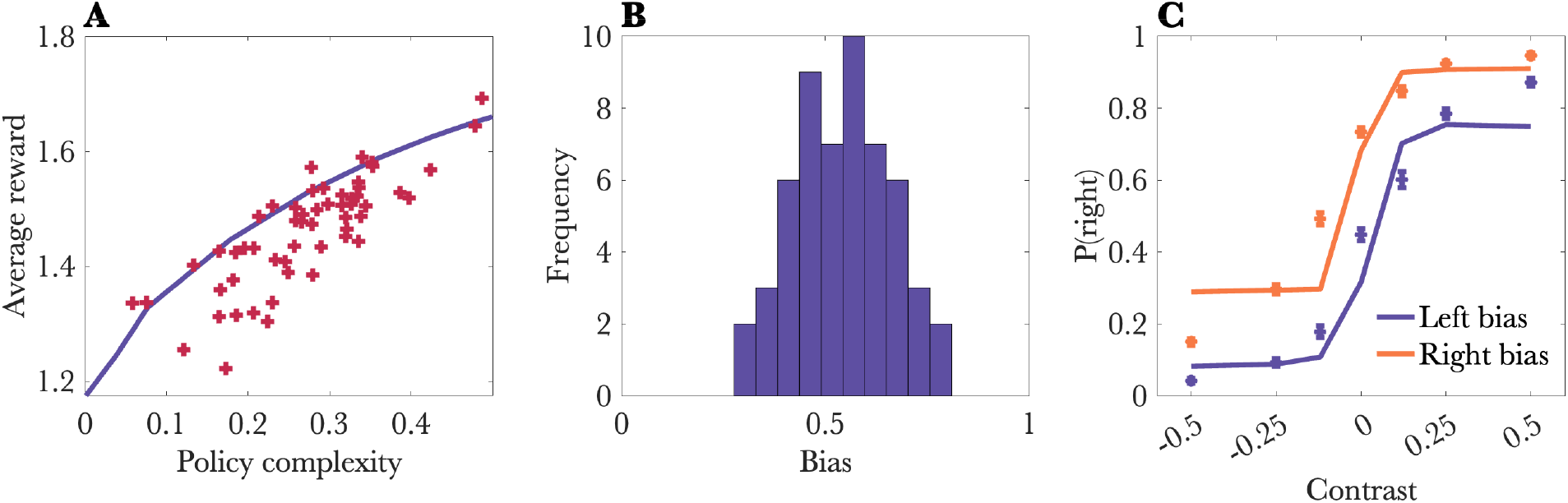
Behavioral results. (A) Task performance was close to the optimal reward-complexity frontier. Each cross represents a single session. Note that a few points are above the curve due to noise in estimation of policy complexity. (B) Histogram of bias (marginal probability of choosing “right”) across sessions. (C) Probability of choosing “right” conditional on stimulus contrast and the session-specific bias. Negative contrast values represent stimuli presented on the left; positive contrast values represent stimuli presented on the right. Solid lines show the model fit. All error bars show standard errors of the mean.

To gain an intuition for what different levels of policy complexity mean behaviorally, we can focus on two aspects of behavior: perseveration and stochasticity. Animals with low policy complexity are perseverative, choosing actions with high marginal probability. In other words, animals will tend to continue choosing an action that may no longer be relevant on the current trial. Animals with low policy complexity are also more stochastic in their action choices. Together, perseveration and stochasticity reduce the average reward for low-complexity policies.

We next tested whether the functional form of the optimal policy (see Materials and Methods) fit the choice data well. The optimal policy has only a single free parameter (the inverse temperature) which controls the balance between reward maximization and policy compression. When this parameter is small, the psychometric function should be shifted in the direction of high frequency actions, which we estimated using the session-specific bias (Fig. 3B). The optimal policy fit the data well, exhibiting a pronounced shift in the psychometric function depending on the session-specific bias (Fig. 3C). Removing the bias term from the policy increased the Bayesian Information Criterion (ΔBIC = 378), thus supporting its inclusion.

Having established the behavioral plausibility of policy compression, we turned to an analysis of dopamine responses at the time of outcome. Linear regression with outcome, value, and policy cost regressors (see Materials and Methods) revealed significant effects for all three (Fig. 4A).

**Figure 4:**
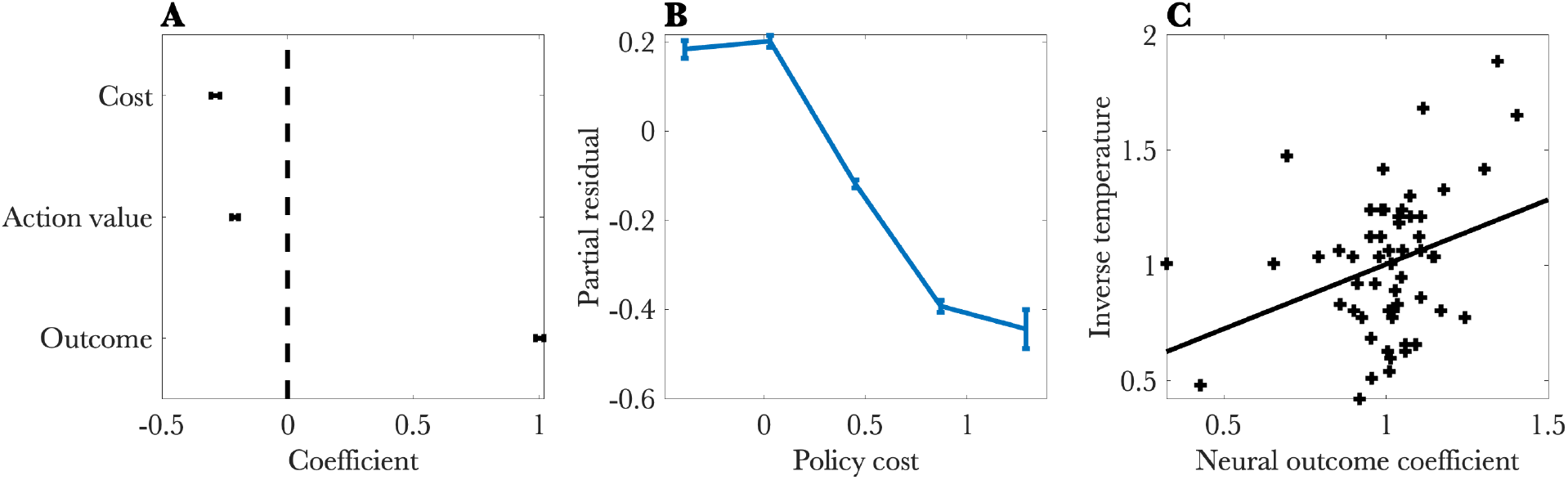
Neural results. (A) Regression coefficients for a linear model predicting the dopamine response at the time of reward feedback. (B) Partial residual plot for the policy cost regressor. (C) Behaviorally estimated inverse temperature plotted against the coefficient for the outcome regressor. Each cross represents a single session. All error bars show standard errors of the mean.

Consistent with a reward prediction error, the outcome effect was positive (*t* = 62.838, *p <* 0.0001), and the value effect was negative (*t* = −15.723, *p <* 0.0001). Critically, the cost effect was negative (*t* = −15.476, *p <* 0.0001; Fig. 4B), consistent with the hypothesis that dopamine signals drive reinforcement learning away from high complexity policies. Removing the cost term from the model increased the Bayesian Information Criterion (ΔBIC = 228). Note that cost and outcome are correlated (*r* = 0.53), so the model comparison result is important for supporting our claim that the cost term is explaining substantial additional variance beyond its shared variance with the outcomes.

Finally, we tested whether the neural and behavioral results align with each other. According to some models (Lai and Gershman, 2021, 2024), the inverse temperature controls both the reward-compression trade-off and the degree of reward esnsitivity in the reward prediction error. This implies that the outcome coefficient in the neural regression should correlate with the inverse temperature fit to behavior, consistent with the experimental data (*r* = 0.33, *p <* 0.02; Fig. 4C).

## Discussion

Our study provides behavioral and neural evidence for policy compression in mice performing a perceptual decision task. Behaviorally, mice approximate the optimal reward-compression frontier, producing patterns of bias quantitatively consistent with the capacity-limited optimal policy. Neurally, dopamine responses to reward outcomes were suppressed by policy complexity, consistent with reinforcement learning models of policy compression (Lai and Gershman, 2021, 2024), and in contrast to models that locate capacity limits outside of the brain’s error-driven reinforcement learning system (Collins et al., 2017).

The idea that an information bottleneck constrains dopamine is buttressed by prior work. Schütt et al. (2024) showed that the population of dopamine neurons forms an efficient code for reward, with tuning curves that maximize information rate subject to a constraint on firing rate. At slower timescales, dopamine may also itself control bottlenecks by modulating sensitivity to sensory and reward signals (FitzGerald et al., 2015; Mikhael et al., 2021; Bari and Gershman, 2023), and by calibrating cognitive effort (Westbrook and Braver, 2016). In this study, we only examined the fast timescale component (phasic responses to reward).

We still lack a biologically plausible circuit model that synthesizes all of these observations. Our data suggest that any such circuit model should include projections from policy-sensitive regions to midbrain dopamine neurons. Searching for such projections will need to start with the identification of regions computing policy cost.

## Acknowledgments

Shuze Liu and Bilal Bari provided helpful comments on an earlier draft. SG is supported by NIH grant U19 NS113201-01 and Air Force Office of Scientific Research grant FA9550-20-1-0413. AL is supported by grant 213465 from the Wellcome Trust.

## Code and data availability

All code and data for reproducing the analyses reported in this paper are available at https://github.com/sjgershm/dopamine-complexity.

## Notes

### Competing Interest Statement

The authors have declared no competing interest.

## References

Bari, B. A. and Gershman, S. J. (2023). Undermatching is a consequence of policy compression. Journal of Neuroscience, 43:447–457.

Collins, A. G., Ciullo, B., Frank, M. J., and Badre, D. (2017). Working memory load strengthens reward prediction errors. Journal of Neuroscience, 37:4332–4342.

Eshel, N., Bukwich, M., Rao, V., Hemmelder, V., Tian, J., and Uchida, N. (2015). Arithmetic and local circuitry underlying dopamine prediction errors. Nature, 525:243–246.

FitzGerald, T. H., Dolan, R. J., and Friston, K. (2015). Dopamine, reward learning, and active inference. Frontiers in Computational Neuroscience, 9:166836.

Gershman, S. J. (2020). Origin of perseveration in the trade-off between reward and complexity. Cognition, 204:104394.

Gershman, S. J., Assad, J. A., Datta, S. R., Linderman, S. W., Sabatini, B. L., Uchida, N., and Wilbrecht, L. (2024). Explaining dopamine through prediction errors and beyond. Nature Neuroscience, pages 1–11.

Lai, L. and Gershman, S. J. (2021). Policy compression: An information bottleneck in action selection. In Psychology of Learning and Motivation, volume 74, pages 195–232. Elsevier.

Lai, L. and Gershman, S. J. (2024). Human decision making balances reward maximization and policy compression. PLOS Computational Biology, 20:e1012057.

Lak, A., Okun, M., Moss, M. M., Gurnani, H., Farrell, K., Wells, M. J., Reddy, C. B., Kepecs, A., Harris, K. D., and Carandini, M. (2020). Dopaminergic and prefrontal basis of learning from sensory confidence and reward value. Neuron, 105:700–711.

Larsen, W. A. and McCleary, S. J. (1972). The use of partial residual plots in regression analysis. Technometrics, 14:781–790.

Mikhael, J. G., Lai, L., and Gershman, S. J. (2021). Rational inattention and tonic dopamine. PLoS Computational Biology, 17:e1008659.

Schultz, W., Dayan, P., and Montague, P. R. (1997). A neural substrate of prediction and reward. Science, 275:1593–1599.

Schütt, H. H., Kim, D., and Ma, W. J. (2024). Reward prediction error neurons implement an efficient code for reward. Nature Neuroscience, pages 1–7.

Tishby, N. and Polani, D. (2010). Information theory of decisions and actions. In Perception-action cycle: Models, architectures, and hardware, pages 601–636. Springer.

Westbrook, A. and Braver, T. S. (2016). Dopamine does double duty in motivating cognitive effort. Neuron, 89:695–710.

